# Induction of territorial behavior and dominance hierarchies in laboratory mice

**DOI:** 10.1101/2024.06.19.599689

**Authors:** Dorian Battivelli, Lucas Boldrini, Mohit Jaiswal, Pradnya Patil, Sofia Torchia, Elizabeth Engelen, Luca Spagnoletti, Sarah Kaspar, Cornelius T. Gross

**Affiliations:** Epigenetics & Neurobiology Unit, EMBL Rome, European Molecular Biology Laboratory, Via Ramarini 32, 00015 Monterotondo (RM), Italy

## Abstract

Territorial behaviors comprise a set of coordinated actions and response patterns found across animal species that promote the exclusive access to resources. House mice are highly territorial with a subset of males consistently attacking and chasing competing males to expel them from their territories and performing urine marking behaviors to signal the extent of their territories. Natural variation in territorial behaviors within a mouse colony leads to the formation of dominance hierarchies in which subordinate males can reside within the territory of a dominant male. While the full repertoire of such territorial behaviors and hierarchies has been extensively studied in wild-derived mice in semi-natural enclosures, so far they have not been established in the smaller enclosures and with the genetically-defined laboratory strains required for the application of neural recording and manipulation methods. Here, we present a protocol to induce an extensive repertoire of territorial behaviors in small enclosures in laboratory mice, including a method for the simultaneous tracking of urine marking behavior in mouse pairs. Using this protocol we describe the emergence of robust dominant-subordinate hierarchies between pairs of CD1 outbred or CD1xB6 F1 hybrid mice, but unexpectedly not in C57BL/6 inbred animals. Our behavioral paradigm opens the door for neurocircuit studies of territorial behaviors and social hierarchy in the laboratory.

## Introduction

Social aggression and defense are core adaptive behaviors of organisms living together in hierarchical or territorial communities^1, 2^. Such territorial behaviors have been best described in animal populations in the wild, but isolated aspects of social aggression and defense can be elicited in the laboratory and neuroscientists have elaborated a rudimentary understanding of the neuroanatomy and pharmacology of these behaviors. However, while much speculation has been aimed at drawing links between territorial behaviors in animals and various types of social aggression, violence, and associated mental states and disorders in humans, we still know too little about the neurobiological basis of these behaviors to draw mechanistic parallels^3–9^. One impediment to the neuroscientific study of territorial behaviors derives from the difficulty in applying the invasive neural recording and manipulation techniques necessary to understand behaviors at the circuit level to populations of animals in the wild or semi-natural environments required to elicit the full repertoire and dynamics of territorial aggression, defense, and dominance behaviors.

Much of our knowledge about the ethology of competitive behaviors derives from the study of the house mouse (*Mus musculus*). When observed in the wild or in large semi-natural environments, populations of house mice form robust social structures characterized by the formation and maintenance of exclusive and semi-exclusive territories by dominant males and the selective mating behavior of females across these territories^10–19^. These structures channel and limit competition among individuals that we can assume provides for an adaptive distribution of natural and reproductive resources. The territorial behavioral repertoire consists of attack and chase behavior elicited by the intrusion of an unfamiliar male into the territory of a dominant animal and the systematic patrolling and urine marking of territorial boundaries by the dominant animal. Dominant males make furtive incursions into the territories of other males, but typically show escape behaviors when approached by the dominant of adjacent territories. Males who reside within the territories of dominant animals adopt subordinate behaviors, including defensive upright postures and hiding which signal submission and help avoid conflict. While high levels of aggression are seen when animals establish territories or when those territories are disturbed, conflict typically subsides when the population is at steady state^12, 13, 20^.

In the laboratory, on the other hand, neurobiologists have primarily used the resident-intruder test to study social aggression in mice. In this test an unfamiliar male is introduced into the home cage of the experimental animal which then proceeds to repeatedly attack the intruder. A variety of lesion, pharmacology, cFos-dependent neural activity mapping, and neural recording and manipulation techniques have been applied to the resident intruder test to identify brain structures, neuromodulators, and circuits that control social aggression in mice^2, 21, 22^. However, the limited space available to mice in the resident-intruder test as well as the familiarity advantage bestowed on the resident does not allow for the expression of the full repertoire of chase, escape, hiding, and marking that characterizes territorial behavior and the evolution of stable dominance hierarchies in semi-natural environments^23–26^.

Here we describe the establishment and validation of a novel laboratory apparatus that elicits the full repertoire of territorial behaviors and allows for the establishment of robust dominance-subordinate hierarchies between male mice. Critically, our apparatus is sufficiently small to be amenable to tethered recording and manipulation techniques required by current neural circuit approaches and elicits the emergence of stable dominant-subordinate hierarchies within hours, a time frame compatible with *in vivo* neural recording methods. We benchmarked the apparatus using CD1 outbred mice, a strain that exhibits robust social aggression in the resident-intruder test^27–29^. However, we found that C57BL/6 inbred mice did not evolve stable dominant-subordinate hierarchies despite showing significant levels of social aggression. In contrast, animals resulting from a cross between CD1 and C57BL/6 mice (F1 hybrid, CD1xB6) succeeded in establishing robust hierarchies, offering an avenue for the application of transgenic mouse lines necessary for cell-type specific neural recording and manipulation to the study of territorial behaviors and dominance hierarchies. Finally, we used our apparatus to develop and validate a protocol for the simultaneous, dual-color quantification of urine marking behavior in dominant-subordinate pairs.

## Results

### Establishment of testing apparatus

In order to allow the expression of a wide repertoire of territorial behaviors between two competing male laboratory mice, we built an experimental device consisting of two large compartments (120 x 60 cm, 70 cm height) connected by a removable barrier (4 x 70 cm; **Figure 1A**). We chose to validate our setup using outbred CD1 mice because this strain has been shown to express robust resident-intruder aggression in laboratory tests^28, 29^. Initially, one of a pair of unfamiliar sexually experienced adult males was placed into each of the two compartments of the apparatus and monitored for 48 hours. Following this habituation period, the barrier was opened and the males were allowed to interact for a period of 2 hours (**Figure 1A**). Behaviors were recorded by a pair of overheard cameras during the first and last 20 minutes of the territorial challenge period (**Figure 1B**). Manual scoring from video was used to quantify and track the timing of social behaviors (attack, chase, flight, defensive upright posture; **Figure 1C**), while two-animal automated tracking from video was used to extract the animals x-y position and calculate time spent in a set of predefined regions of interest within the apparatus (hiding, exploration, opponent’s resources investigation and total locomotion; **Figure 1B**).

**Figure 1.**
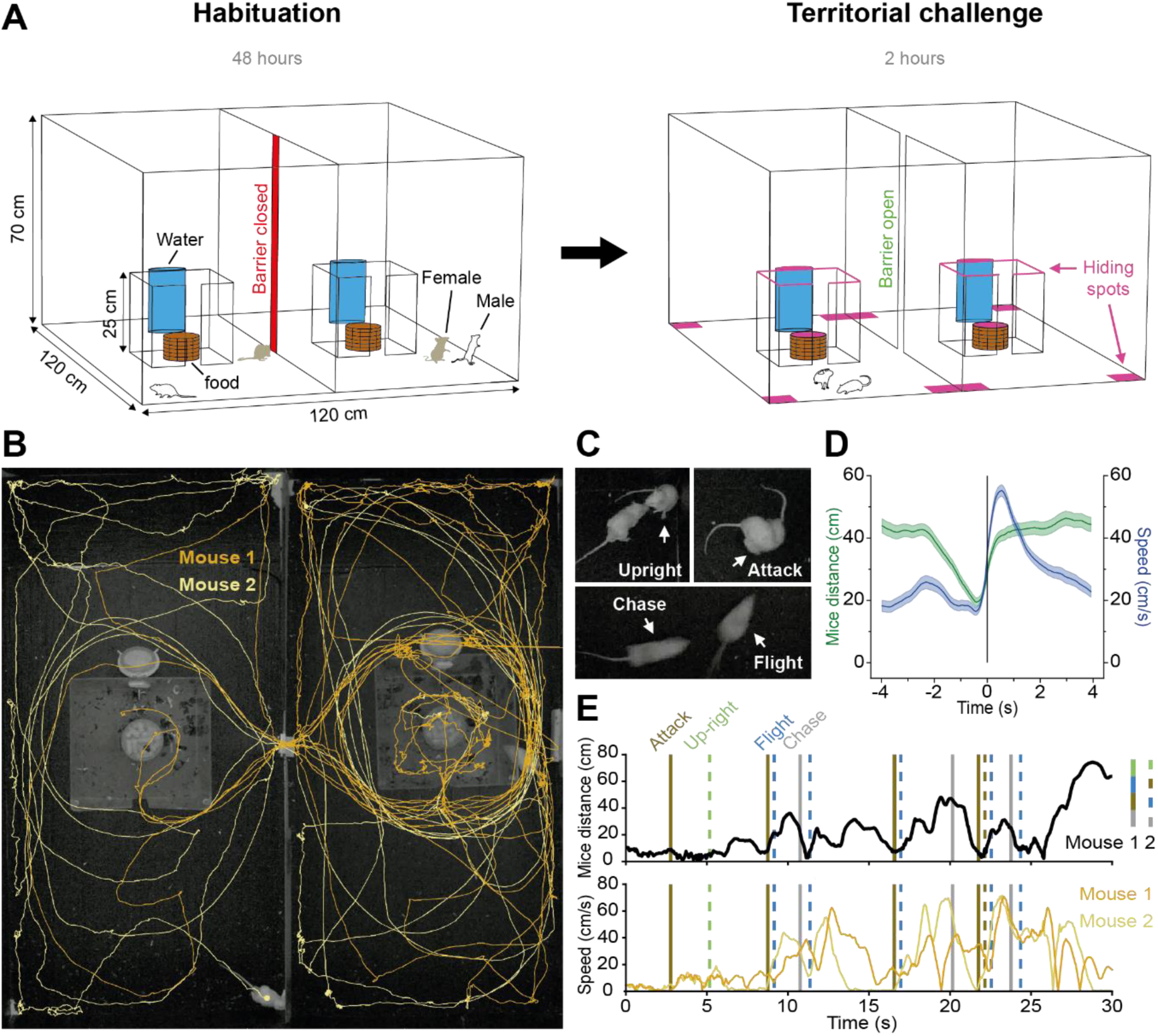
Design of semi-natural testing apparatus for the study of territoriality in laboratory mice. (**A**) Unfamiliar CD1 adult males (white) mated for one week with females (brown) were habituated for two days in large adjacent environments connected by a removable gate and with free access to a central shelter with food and water. On the third day, females were removed and the gate was opened for 2 hours to allow the two males to explore the full apparatus and display territorial behaviors that were recorded by overhead video cameras. (**B**) Representative path plots of two interacting male mice extracted from video recordings (10 minutes). (**C**) Representative video frames showing selected territorial behaviors manually scored from video recordings (upright posture, attack, chase, flight). (**D**) Quantification of flight behavior showing speed of fleeing animal and distance between the two animals during the late interaction phase (mean ± SEM, t = 0 indicates flight onset; N = 176 flights, N = 7 mice). (**E**) Representative 30 sec traces of distance between two mice (top) and speed (bottom) with territorial behaviors for each mouse indicated by vertical lines (attack, upright posture, flight, chase).

Mice pairs consistently showed a stereotyped progression of territorial behaviors following the opening of the barrier. Initially, both mice explored the apparatus, followed by a period in which both mice engaged in repeated bouts of attack and disengagement. Attacks were interspersed by bouts in which one of the two males chased his opponent and the other showed high speed flight, a behavior that typically peaked within one second after onset and reached speeds of over fifty centimeters per second (**Figure 1D**). Flights were often punctuated by the fleeing animal taking a defensive upright posture in which the animal reared on its high limbs and faced the aggressor. Eventually, attacking behavior became more infrequent and one of the two mice consistently showed flight responses when approached by the other mouse (**Figure 1E**), a behavioral pattern associated with increased immobility and hiding behavior in which the animal remained in the corners or along the walls of the compartments or sitting on top of the water bottle or plexiglass home cage for long periods of time. These observations suggested that the behavior of the two males gradually diverged during the two-hour observation period and pointed towards the emergence of a social hierarchy.

### Emergence of dominance hierarchies

We first sought to quantify the evolution of the relationship between mouse pairs. Consistent with an initial balance in territorial behaviors between opponents we found that there was no significant deviation from parity in the first twenty minutes of the challenge period for attack, chase, flight, upright posture, hiding, exploring, and locomotion (**Figure 2A**). To follow the subsequent evolution of hierarchical behaviors we calculated difference scores for each behavior between early (0-20 minutes) and late (100-110 minutes) phases of social interaction. Differences in the number of attacks, chases, and flights, and time spent hiding increased significantly over time, while those for exploration, opponent’s resources investigation, locomotion and upright postures did not change significantly (**Figure 2A**), although the latter showed a trend for decreasing over time, presumably as the overall number of attacks and chases decreased and flights without chase became the predominant defensive strategy. These findings demonstrate that the territorial behaviors of mouse pairs diverged significantly over time and suggest the emergence of a social hierarchy.

**Figure 2.**
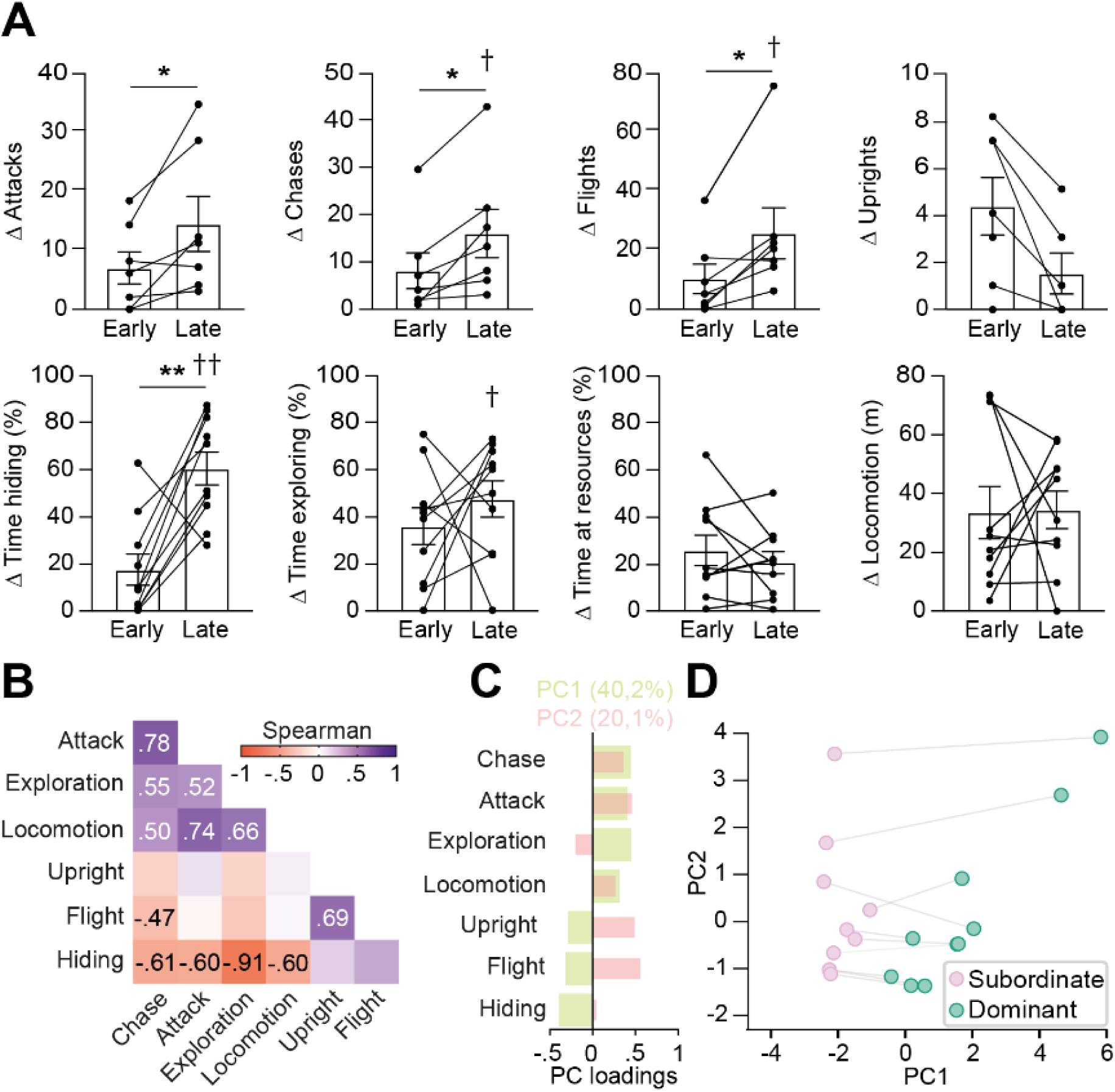
CD1 mice display robust territorial behaviors. (**A**) Quantification of the absolute value of differences in territorial behaviors between mice pairs (Attacks, chases, flights, upright posture: N = 7 pairs, 3 pairs were excluded because one mouse hid in a location inaccessible to its opponent during the entire late interaction period; hiding, exploring, opponent’s resources investigation, locomotion: N = 10 pairs; Wilcoxon matched-pairs signed rank test: *P < .05, **P < .01, ***P < .001; permutation test to evaluate differences between opponents within each time period: †P < .05, ††P < .01; mean ± SEM). (**B**) Correlation matrix indicating how behaviors covary within the mouse population during the late interaction phase (N = 20 mice; only significant correlations are indicated). (**C**) Individual loadings of the first two principal components (PC1: variance = 40,2%, PC2: variance = 20,1%) carried out on territorial behaviors (chase, attack, exploration, locomotion, upright posture, flight, hiding). (**D**) Plot of PC1 and PC2 values for each pair, with the mouse with the higher PC1 value in each pair labeled as dominant and the other as subordinate.

To understand how individual behaviors might be coordinated to reflect a coherent territorial strategy we performed a within-animal correlation analysis of behavioral measures across the entire CD1 population during the late phase of the observation period. Two groups of behaviors emerged that showed positive within-group and negative between-group correlations (**Figure 2B**). The first group comprised attack, chase, exploration, and locomotion and the second included flight, hiding, and, to a lesser extent, upright postures. These correlations demonstrated that the more aggressive a mouse is, the more it explores its environment and the less likely it is to flee or hide. Because dominant wild-derived mice in semi-natural enclosures are more aggressive and explorative and spend less time hiding than subordinate animals^12, 30^ the behavioral organization observed suggested that both hierarchy and territory were firmly established at the conclusion of the two hour challenge period.

Next, we carried out a principal component analysis (PCA) of all behaviors to identify optimal behavioral factors that might reliably describe the global behavioral patterns of each animal in the pair (**Figure 2CD**). The first principal component (PC1) accounted for approximately 40% of the total variance in behavior and was strongly positively correlated with attack, chase, exploration, and locomotion and negatively correlated with flight, upright postures, and hiding (**Figure 2C**). The second principal component (PC2), on the other hand, explained about 20% of the variance in behavior and was positively correlated with measures of social engagement (attack, chase, locomotion, flight, upright posture) and negatively correlated with non-social behaviors (exploration). The PCA indicates that behavior in our test can be described by two major orthogonal factors, one that reflects dominance status and the other that reflects social engagement. Finally, we plotted each mouse pair by their PC1 (dominance) and PC2 (social engagement) scores and labeled each mouse as dominant or subordinate based on the relative magnitude of their PC1 score (**Figure 2D**). Taken together, these results suggest that within a two hour period our apparatus is able to elicit the emergence of robustly divergent territorial behavior strategies that reflect the establishment of a stable social hierarchy between pairs of laboratory mice.

### Strain comparison

Next, we set out to determine whether similar social hierarchies could be elicited in the C57BL/6 mouse strain. This strain is widely used in behavioral neuroscience studies due to the availability of genetically modified congenic lines that aid researchers in carrying out cell-type specific neural monitoring and manipulation. Consistent with previous studies that reported a low penetrance of resident-intruder aggression in C57BL/6 mice^27^, we found that, while all CD1 pairs exhibited attack behaviors during the two hour observation period, only 61% (11/18) of C57BL/6 pairs showed attack (**Figure 3A**), although C57BL/6 mice did exhibit significant exploratory behavior during this time confirming that the lack of aggression was not secondary to an absence of social interaction (**Figure 3B**; **Figure S2**). However, no significant differences in territorial behaviors emerged during the two hour period in C57BL/6 pairs (**Figure S2**), a finding that persisted even in the subset of mice pairs that exhibited aggression (**Figure S3**), suggesting that this strain was not able to develop robust social hierarchies. Moreover, while time spent in physical proximity decreased significantly in CD1 pairs between the early and late periods, proximity remained high throughout the test in C57BL/6 mice (**Figure 3A**) showing that the failure to develop a hierarchy was linked to a failure to socially disengage. In line with these observations, both dominance (PC1) and social engagement (PC2) scores in C57BL/6 mice clustered closely together and the difference in dominance scores in each pair was significantly smaller than in CD1 mice (Kruskal-Wallis test, H = 19, ***P < .001; **Figure 3CD**). Furthermore, behavioral measures were poorly correlated in C57BL/6 mice and in some cases showed anomalous correlations (e.g. positive correlation between attack, upright posture, and flight; **Figure 3E**). Together, these data point to a disorganization of territorial behavior in the C57BL/6 strain in our test.

**Figure 3.**
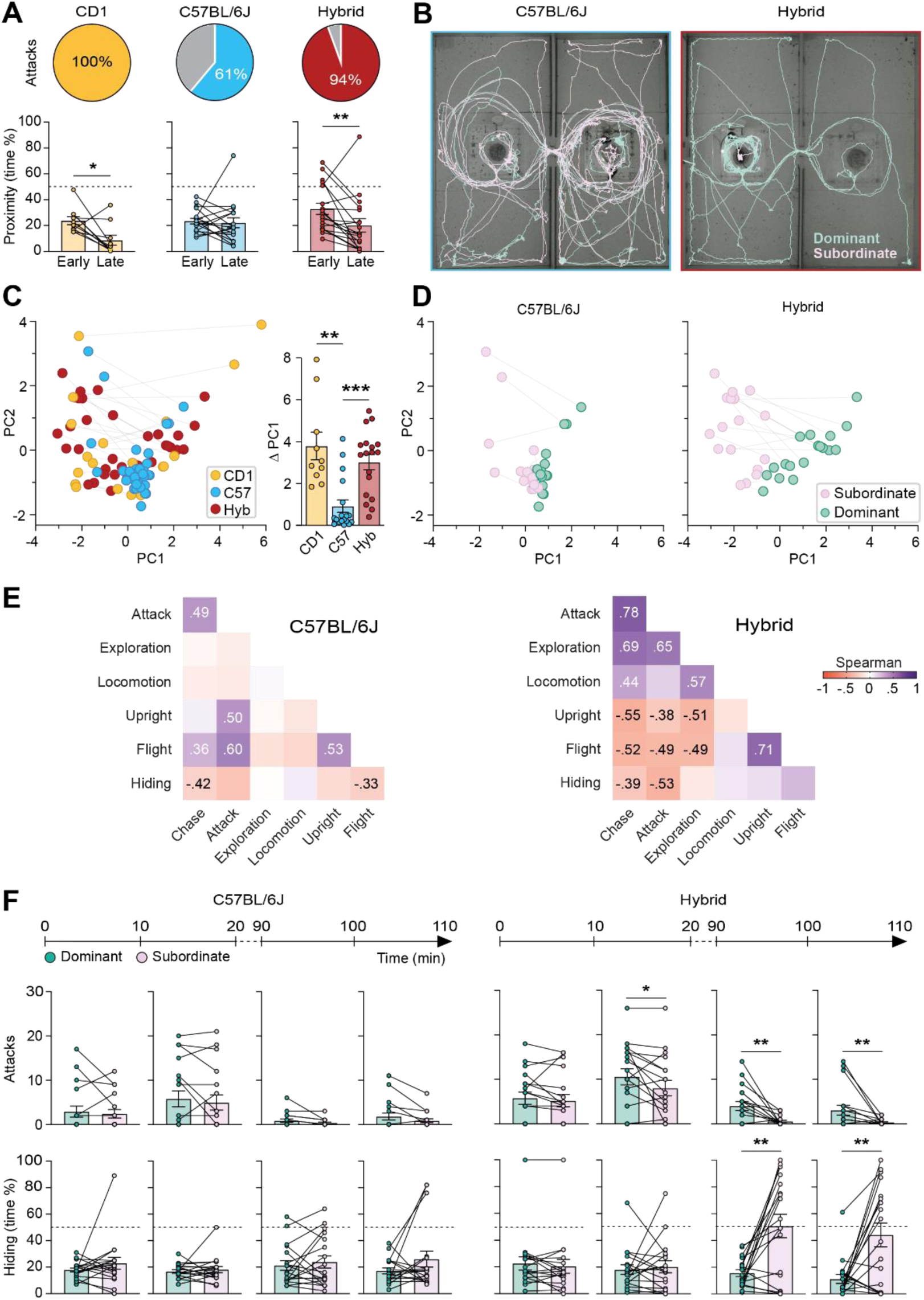
Poor territoriality in C57BL/6, but not CD1xB6 F1 hybrid mice. (**A**) Percentage of mice pairs showing attack behavior for each genetic strain (top). Quantification of proximity between mice within each pair (bottom; CD1: N = 10 pairs, C57BL/6 and hybrids: N = 18 pairs; Wilcoxon matched-pairs signed rank test: *P < .05; **P < .01; ***P < .001; mean ± SEM). (**B**) Representative path plots of two interacting male mice (dominant, subordinate) extracted from video recordings (10 minutes from the late phase) for a pair of C57BL/6 (left) and CD1xB6 F1 hybrid (right) mice. (**C**) Plot of PC1 and PC2 values for all mice pairs (left) labeled by strain. Comparison of PC1 difference (dominant minus subordinate) values between strains revealed a significant difference between C57BL/6 mice and the other strains (right; N = 18 pairs for each strain; Dunn correction for pairwise comparisons, **P < .01, ***P < .001; mean ± SEM). (**D**) Plot of PC1 and PC2 values for C57BL/6 (left) and CD1xB6 F1 hybrid (right) mice pairs labeled as dominant or subordinate according to PC1 score. (**E**) Correlation matrices between territorial behaviors for C57BL/6 (left) and CD1xB6 F1 hybrid (right) populations during the late interaction phase (N = 36 mice; only significant correlations are indicated). (**F**) Quantification of the evolution of behavioral differences between dominant and subordinate C57BL/6 (left) and CD1xB6 F1 hybrid (right) mice across subintervals (10 minutes) of the early and late observation periods (N = 18 pairs for each strain; Wilcoxon matched-pairs signed rank test; *P < .05, **P < .01, ***P < .001; mean ± SEM).

Finally, we examined whether hybrid mice that result from the crossing of CD1 outbred and C57BL/6 inbred mice (F1 hybrid, CD1xB6) could develop robust social hierarchies in our test. If successful, hybrid mice could permit the use of heterozygous Cre-driver alleles deriving from the C57BL/6 parent that would otherwise be cumbersome to derive on or backcross to the CD1 strain. Fortunately, hybrid mice showed a similar evolution and organization of territorial behavior and social hierarchy as that found in CD1 mice (**Figure 3A-E**). In particular, 94% of hybrid mice pairs showed aggression (**Figure 3A**) and hybrid mice showed significant exploratory behavior (**Figure 3B**), a significant decrease in proximity behavior (**Figure 3A**), and the emergence of robust differences in dominance (PC1; **Figure 3CD**), and a pattern of correlations between behavioral measures similar to that seen in CD1 mice (**Figure 3E**).

A more detailed analysis of the evolution of behavioral measures over time confirmed a failure of C57BL/6 pairs to show any significant differences in attack or hiding across the observation periods, while CD1xB6 hybrid mice showed a gradual and robust emergence of hierarchy (**Figure 3F**; **Figure S4**). Notably, both hybrid and CD1 mice pairs showed a sequential emergence of differences in territorial behaviors with differences in attacks preceding differences in hiding (**Figure 3F**; **Figure S1**). The staggered emergence of behavioral differences suggests that defensive hiding may be a direct consequence of the differences in aggression, rather than being a separately developing hierarchical trait. Overall, our results demonstrate that while C57BL/6 mice failed to show the emergence of an organized social hierarchy, CD1xB6 hybrids could do so robustly and in a manner similar to CD1 mice.

### Urine marking behavior

Urine marking plays a key role in the signaling of territorial boundaries in mice^31^. However, until now, urine marking quantification methods were restricted to testing one animal at a time, making it difficult to extract relative territorial behavior measures. To simultaneously quantify urine marks for both animals in our test we developed a dual-dye labeling protocol based on the systemic delivery of fluorescein and erythrosin b dyes to mouse pairs and the image-based quantification of these dyes in urine excreted during exploration of the apparatus. Following the successful establishment of hierarchies, mice were briefly removed from the apparatus, injected with dyes, returned to their home cage for 45 minutes, and then returned to the apparatus and allowed to explore freely for 60 minutes. Upon completion of the test, the floor of the apparatus was imaged with a color camera and patches of the two excreted dyes segmented and quantified (**Figure 4AB**). A quantification of the extent of dye marks revealed a significant difference in marking behavior between CD1xB6, but not B6 mice pairs (**Figure 4B**). Among hybrid, but not B6 pairs, mice with higher dominance (PC1) scores showed significantly more extensive marking than subordinate mice (**Figure 4C**). Such mice also consistently marked across the entire apparatus, consistent with their showing territorial behavior across both compartments. Finally, we tested whether the dyes adversely affected mouse behavior at the concentrations used. No significant differences in total locomotion or anxiety as measured in the elevated plus maze (Kruskal-Wallis test – acute locomotion: H = 0.30, P = .83; chronic locomotion, H = 1.19, P = .55; acute anxiety: H = 0.38, P = .83; chronic anxiety, H = 0.25, P = .88), body weight (two-way repeated measure ANOVA – main effect of time: F_4.07, 85.5_ = 1.53, P = 0.20; main effect of treatment F_2, 21_ = 1.17, p = .33); time x treatment interaction: F_14,15_ = 1.07, P = 0.39), or olfactory preference for dye-containing urine were detected following either one or six daily injections (**Figure S5A-D**). These results establish a method for the quantification of urine marking in interacting mouse pairs and confirm that the behavioral hierarchies that emerge in our apparatus also extend to territorial marking behaviors.

**Figure 4.**
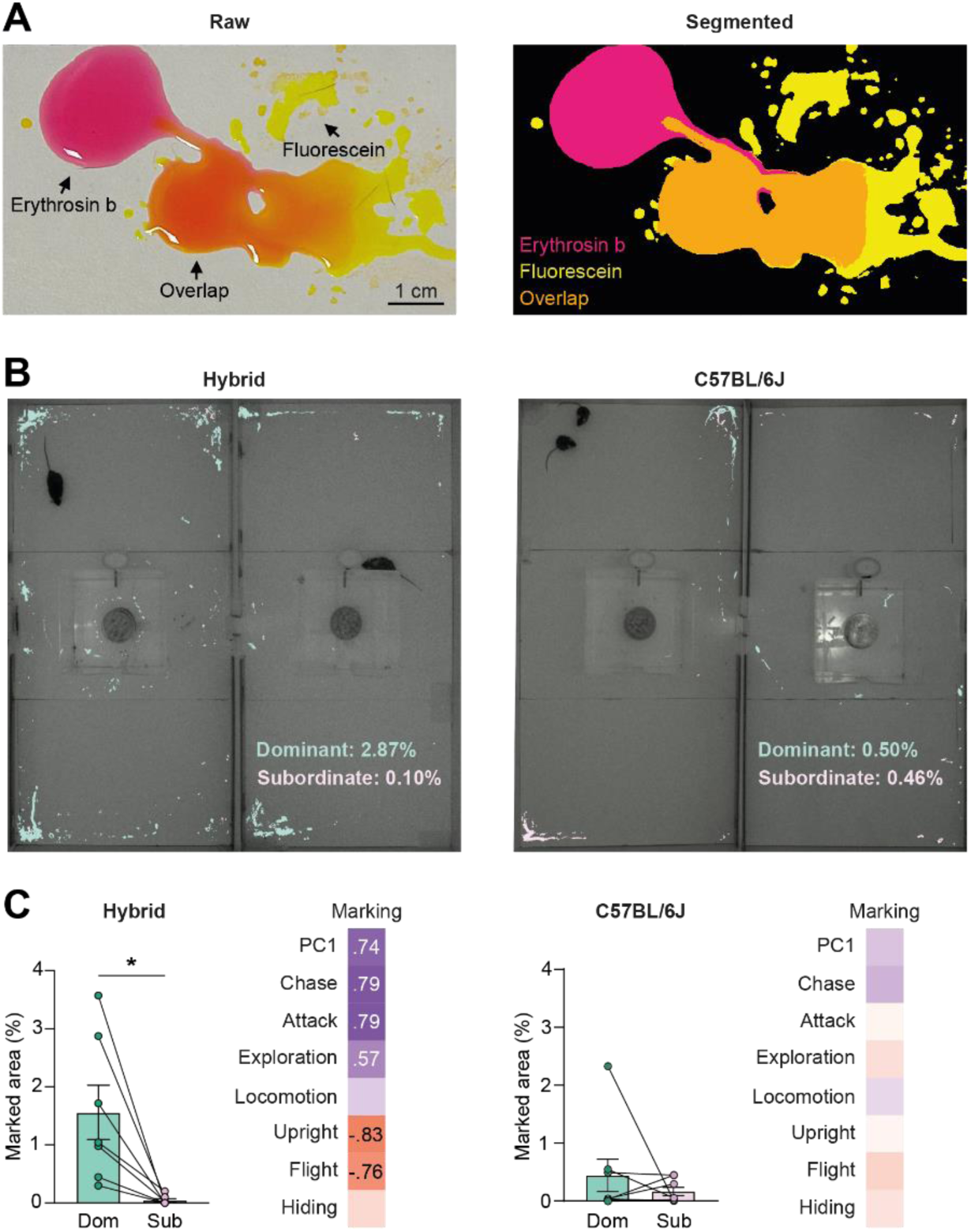
Simultaneous quantification of urine marking in interacting mice. (**A**) Representative urine marks deposited by a mouse injected with erythrosin b (fuchsia) or fluorescein (yellow). (**B**) Representative images of cumulative urine marking of two interacting mice of the C57BL/6 (left) or CD1xB6 F1 hybrid (right) strain (10 minutes). (**C**) Quantification of percentage of marked area for pairs of dominant and subordinate mice for the C57BL/6 (left) or CD1xB6 F1 hybrid (right) strains. Correlation matrices indicate the association between urine marking behavior and other territorial behaviors within the C57BL/6 (left) or CD1xB6 F1 hybrid (right) population (C57BL/6: N = 8 pairs, hybrids: N = 7 pairs; only significant correlations are reported; Wilcoxon matched-pairs signed rank test: *P < .05; **P < .01; ***P < .001; mean ± SEM).

## Discussion

We report the development and validation of a novel behavioral testing apparatus to study territorial behaviors and hierarchies in laboratory mouse strains. Our work aimed to fill a gap between the widely used resident-intruder test for measuring social aggression in the laboratory and ethological studies describing a wider repertoire of territorial behaviors in large natural and semi-natural enclosures. Although the resident-intruder test can be used to quantify aggressiveness, it fails to elicit the full repertoire of territorial behaviors, including attack, chase, defensive upright, social avoidance, and marking behaviors observed in natural settings. On the other hand, studies in larger semi-natural enclosures are poorly adapted to the application of modern neuroscience methods that typically require tethering the animals to electronic devices for recording and stimulation. In addition, doubts have been raised about the use of laboratory inbred strains for studying aggression-related behaviors in mice as they show reduced aggression relative to wild-derived strains^27, 32, 33^. Here, we found that by using a somewhat larger apparatus (120 cm x 120 cm) we could reliably elicit the full repertoire of territorial behaviors in the laboratory, including the establishment of social hierarchies described in semi-natural and natural enclosures. An apparatus of this size is readily adapted to tethered recording and manipulation methods in mice and suggests that our test offers a means to study the full repertoire of territorial behaviors at a neurocircuit level.

A major finding of our study is that C57BL/6 inbred mice were not able to establish robust dominance relationships. Indeed, even under conditions that were intended to promote social competition (i.e. sexual experience, extensive habituation to testing apparatus) we observed that almost half of inbred mice did not show aggressive behavior. While the reduced penetrance of aggression has been well documented in inbred mice^27, 32, 33^, we found that even among C57BL/6 mice pairs that did show aggression, the relationship did not evolve into a stable hierarchy over time as reflected in differences in hiding, exploration, and marking behaviors as seen in CD1 animals (**Figure 3F**, **S3A-B**). Importantly, the lack of hierarchy in C57BL/6 mice was also reflected in a failure of these mouse pairs to avoid each other as the encounter progressed (**Figure 3A**), suggesting that they lacked the capacity to socially disengage under these circumstances. A persistence in affiliative social behaviors, including communal sleeping, has been previously associated with the inability to form social hierarchies in mice^32^. Our failure to find robust hierarchy formation in C57BL/6 mice in our apparatus appears to contradict a recent study which reported territory formation in this strain in semi-natural field enclosures^34^. However, in this study, the measure of territoriality was limited to space occupancy patterns without providing an indication of the quality of interactions between opponents. Conversely, previous observations showed that pairs of unfamiliar male C57BL/6 mice released for 24 hours in a complex environment showed reduced aggression and consistently formed communal nests^32^, supporting our observations in this strain.

In contrast, we found that CD1xB6 F1 hybrid mice were able to express territorial behaviors and establish social hierarchies in a manner similar to CD1 outbred mice. This strain offers the possibility of accessing transgenic alleles commonly maintained on the C57BL/6 strain and thus could facilitate the use of Cre recombinase driver lines that allow cell-type specific neural monitoring and manipulation, or other reporter alleles maintained in a heterozygous state. Nevertheless, some differences in territorial behavior were detected between CD1xB6 hybrids and the CD1 parental strain. Subordinate CD1 mice spent the majority of time in highly inaccessible areas of the apparatus, such as on top of the water bottle or home enclosure walls, while this was seen only transiently in hybrid mice. This discrepancy may be related to the somewhat lower intensity of aggression seen in hybrid compared to CD1 mice (14.8 vs 22.8 attacks/20 minutes, respectively; **Figure S1, S4**) or, alternatively, it may indicate a subordinate strategy in this strain less dependent on passive avoidance.

In all CD1 and hybrid mice pairs we studied we observed the emergence of a clear dominance hierarchy, with one animal exhibiting dominance over the other in the entire apparatus. The failure to observe the emergence of stable side-by-side territorial behaviors in which each animal defends and marks its own compartment was unexpected, as some studies have reported the formation of such neighboring territories in wild-derived mice in laboratory enclosures similar to ours^12, 13^. An exploration of factors with the potential to influence levels of territoriality, including sexual experience, single housing, and prior social winning failed to promote the reliable emergence of side-by-side territories, although these did occasionally appear at low frequency (unpublished data). At present we do not have an explanation for why such outcomes are rare. Our device may be too small to satisfy a minimal territory requirement^12, 13, 16, 35^ or larger populations may be needed to help stabilize the formation of smaller territories^11, 13, 16, 36^. Alternatively, side-by-side territories could be stabilized by increasing the complexity of the habitat so as to increase hiding spaces and favor the maintenance of territories^19^.

Historically, detailed descriptions of territorial and hierarchical behaviors were carried out on wild and wild-derived outbred mouse strains observed in their natural habitat or later in semi-natural laboratory environments^10, 11, 12, 13, 14, 15–17, 20, 37^. Increasingly, social hierarchies have been studied in the laboratory in inbred mouse strains at the neuroanatomical and circuit level. The reduced levels of social aggression typical of such strains has led researchers to seek alternate behavioral surrogates of dominance, such as the tube and hot spot tests, mating-associated ultrasound vocalizations, and urine marking tests^38–42^. Our test offers an avenue to study *bona fide* measures of male-male social aggression and dominance behavior in the laboratory in a relatively limited space compatible with modern circuit manipulation and monitoring methods. We hope that this more naturalistic approach will facilitate a circuit understanding of aggression and dominance that takes into consideration the social context of these behaviors, and thus may be more relevant to our understanding of the molecular basis of social threat responding and its translation to human behavior.

## Materials & Methods

### Animals

All experimental procedures involving the use of animals were carried out in accordance with EU Directive 2010/63/EU and under approval of the EMBL Animal Use Committee (IACUC) and Italian Ministry of Health License 2021-04-09 to C.G. Animals were born and maintained in a temperature and humidity controlled environment with food and water provided *ad libitum* and 12 hr/12 hr reverse light-dark cycle (lights on at 9:00 PM). Wild-type C57BL/6J and CD-1 mice were bred in our facility from ancestors acquired from Charles River Laboratories (Calco, Italy). To produce CD1xB6 hybrid mice we crossed CD-1 females with C57BL/6J males. All experimental animals were weaned three weeks after birth and housed with male siblings (3-5 per cage).

### Territorial arena

The experimental setup consisted of a white wooden floor, divided in two side-by-side rectangular compartments of equal dimensions (120 x 60 cm). Each compartment was enclosed by white polypropylene panels (70 cm high). For experiments with CD1 mice we added black polypropylene panels to cover the floor and bottom of the walls (15 cm high) to increase contrast and improve video tracking. A removable door in the center wall dividing the two compartments and provided a passageway (4 cm wide) between compartments. In the center of each compartment we placed a roofless plexiglass home cage (25 cm high) with an entrance (4 cm wide) facing perpendicular to the adjacent compartment. The home cage contained a column-shaped wire mesh enclosure (12 cm high, 10 cm diameter) filled with regular chow food pellets and covered by a plexiglass disk and a water dispenser attached to the middle of the wall facing the entrance.

### Territorial behavior testing

One week before their introduction into the territorial arena, unfamiliar non-sibling males were singly mated into a fresh standard holding cage. During this period, males were shaved with a specific pattern on the back to be identified. In order to accustom the animals to the experimental device and allow them to mark their respective environments, each couple was placed in one of the adjacent compartments (in the morning, at the start of the active phase), keeping the gate connecting the two compartments closed during the following 48 hours. At the start of the third day, the females were removed from the device, and after 10 minutes, the connecting gate was opened to let the two males explore the neighboring compartment and interact with each other for 120 minutes.

### Behavior scoring and quantification

Cameras (Basler acA2040-55uc) mounted above each compartment allowed for continuous remote monitoring of the mice during the whole interaction phase. Behaviors were recorded during the first (early stage) and last (late stage) 20 minutes of the experiment (under red light, at 45 fps using Pylon viewer software). For each recording session, the two resulting videos were merged with Bonsai^43^ to obtain a single video output, which was used for behavioral scoring. Between these two periods, we regularly monitored the animals to ensure that there were no severe injuries requiring interruption of the experiment.

Animals were automatically tracked via DeepLabCut^44^. For each line, we trained and applied a distinct model as follows: 200 frames were automatically extracted from different pairs (using k-means method), and 8 body parts were manually annotated on each animal (nose, right and left ears, middle back, right and left side, base and end of the tail). The model was trained (Resnet_50 network) until an accurate detection plateau was reached (between 117,000 and 200,000 iterations, depending on the strain). After all videos were analyzed, we manually corrected the identity swaps.

Resulting pose estimates were used on SimBA^45^ to quantify locomotion and time spent by each mouse in regions of interest. Preliminary data cleaning based on interpolation and smoothing methods was systematically applied to DeepLabCut output. The ROIs were drawn manually. Before processing ROIs for hiding, exploration and proximity quantification, we discarded periods of <1 second occupancy from the raw SimBA quantifications for each ROI. This was intended to reduce tracking artifacts. Hiding behavior was quantified as the sum of time spent in the 12 following ROIs: the 4 corners of each compartment (8 x 8 cm), on the top of the food columns and on the top of the central cages including the top of the water dispensers. Exploration was quantified as the time spent in the opponent’s compartment, minus the time spent in the hiding zones of this same compartment. Investigation of opponent’s resources was quantified as the time spent in the central cage of the opponent’s compartment, minus the time spent on the top of the food column of this cage. The animal proximity was computed as the total time two mice spent less than 10 cm from each other, within the same ROI. Here we considered the following ROIs: each compartment, each central cage, top of each food column, and top of each central cage.

Fighting behaviors (attack, chase, flight and upright posture) were manually annotated with Solomon Coder (RRID:SCR_016041). Manual annotation was chosen following unsuccessful attempts to obtain sufficiently robust automatic tracking at the time of close interactions, necessary for the identification of complex interactions. Attack was defined as a behavioral sequence initiated by lunges toward the opponent and resulting in physical contacts including biting and tumbling. Chase was defined as a rapid movement in pursuit of the opponent, leading to no physical contact. Flight was defined as a sudden acceleration in the opposite direction of the opponent. Upright postures were identified when the mouse sat erect on its hind paws, with the head raised and the forepaws stretching out, leaving the belly exposed.

The characterization of speed and distance around flights was based on the behavior recorded during the late interaction phase. Specifically, around each flight event of every mouse, we identified the moment of the mouse’s maximum acceleration as estimated by calculating the steepest slope in the mouse speed within a 90-seconds asymmetric window around the flight onset (30 seconds before and 60 seconds after). The frames corresponding to the peak acceleration were used as centers for larger, symmetric 8-second windows. For each mouse, the same set of centers were used to compute windows for both distance and speed traces. Finally, all windows from all mice were overlapped and averaged on a frame-by-frame basis.

To evaluate whether behavioral differences pre-existed between opponents, or conversely, emerged as a consequence of the territorial confrontation, we performed a permutation test at early and late stages of interaction. This approach allowed for a comparison, for each behavior, of the observed absolute average difference to a null distribution for the absolute average difference obtained by randomly pairing mice from the whole CD1 cohort. The P-value was determined as the fraction of permutations for which the null average difference was larger than the observed one (i.e. without permutation).

### Hierarchical rank identification

In order to identify social ranks within each pair we performed a principal component analysis (PCA), a dimensionality reduction technique capable of transforming high-dimensional data into a lower dimensional representation. PCA aims to preserve the variance in the original data by identifying and retaining the principal components, which are linear combinations of the original variables. These components capture the most significant sources of variation, with the first principal component explaining the maximum variance in the data. Subsequent components explain decreasing amounts of variance, and together they account for the entire variance present in the original dataset. PCA loadings for PC1 and PC2 represent the contribution of each original variable to the two principal components.

A single PCA was performed for all mice of the three strains. We included the number of attacks, chases, flights and upright postures, the locomotion expressed in cm, and the proportion of time spent in exploration and hiding for each animal during the late interaction phase. For each behavior, data were normalized including all mice of different strains (z-scores).

### Urine marking

To quantify urine marking in two animals simultaneously we developed a dual-color urine marking method. One mouse was injected with fluorescein (39 mg/kg i.p.; Sigma-Aldrich, CAS-No 518-47-8) and the other with erythrosin b (54 mg/kg s.c.; Sigma-Aldrich, CAS-No 16423-68-0). Empirical testing showed that the dyes become visible in urine marks ∼25 minutes after injection and remain visible for at least 12 hours thereafter. Immediately after the territorial challenge the mouse pairs were removed from the arena and injected each with a different dye (fluorescein or erythrosin b) and placed into a fresh standard holding cage. Forty-five minutes later the animals were placed on white paper to check for the presence of dye in the urine and reintroduced for one hour together into the experimental arena with the central gate open. Following removal of the mice, a photo of each compartment was taken under normal room light and processed with Ilastik^46^ to segment and quantify two-color urine marks. The segmentation algorithm was trained using manually annotated images. Marking for each animal was represented as a percentage of total area of the arena.

### Assessment of behaviors associated with dye treatment

#### Anxiety-like behavior and locomotion in Open Field test

Adult C57BL/6 males were habituated to the experimental room for the 2 days preceding the experiment by being transferred there for 2 hours per day in the morning in their homecage. On the day of the experiment, animals were treated with fluorescein, erythrosin b, or saline upon arrival in the testing room before they were introduced, at least 45 minutes later, into an Open Field test chamber (40 x 40 x 35 cm) for 10 minutes where locomotion was recorded by a camera (Basler acA2040-55uc, 45 fps) placed above the device, under red light. We automatically quantified the time spent in the central square of the Open Field test apparatus (20 x 20 cm) and the time spent in the corners (5 x 5 cm) using DeepLabCut and SimBA. The ratio between these two spatial categories was used as a measure of the anxiety-like behavior of the animal (T[center] - T[corners]) / (T[center] + T[corners]). The experiment was repeated after 6 consecutive days of treatment. This test was performed twice, upon the first day of injection (acute) and after 4 days of injection (chronic).

#### Body Weight

Mice were weighed every afternoon (i.e. ∼5 hours after they were injected) and injected and weighed according to the same schedule on the 8th and 10th day after the first day of treatment.

#### Odor preference test

New adult C57BL/6 singly housed males were habituated to the testing room in their original cages for 1 hour a day for 3 days. On the experimental day, each animal was tested for 10 minutes in a clean cage, without bedding and covered by a perforated plexiglass plate. For each trial, three cotton swabs were freshly soaked with a mixture of urine from 8 mice treated with fluorescein, erythrosin b, or saline (40 μl of urine collected from each mouse) and inserted into syringe caps cut at their ends. The constructs were inserted equidistantly into holes of the plexiglass cover along the length axis, placing the odorous end towards the inside of the cage, 7 cm from the ground. The order of the cotton swabs was randomly shifted between trials. Behavior was videotaped from the side (Panasonic SDR-H100, 30 fps) and the time spent sniffing each olfactory cue was scored manually by an experimenter blind to the condition. A preference index was calculated for each dye compared to saline (T[dye] - T[saline]) / (T[dye] + T[saline]).

Manual annotations and behavioral scoring were made by an experimenter blind to the experimental treatment.

### Data analysis and graphs

Statistical tests were performed using GraphPad Prism version 10.2.3 for Windows. Permutation test was performed using customized R script. PCA was performed using a customized Python script. Graphs were generated with Graphpad Prism and further customized with Adobe Illustrator 2024, version 28.5.

### Data and code availability

Custom code written for this study is made available on GitLab platform https://git.embl.de/grp-gross/induction-of-territorial-behavior-and-dominance-hierarchies-in-laboratory-mice. Behavioral and imaging data will be made available upon reasonable request.

## Supplementary Figures

**Figure S1.**
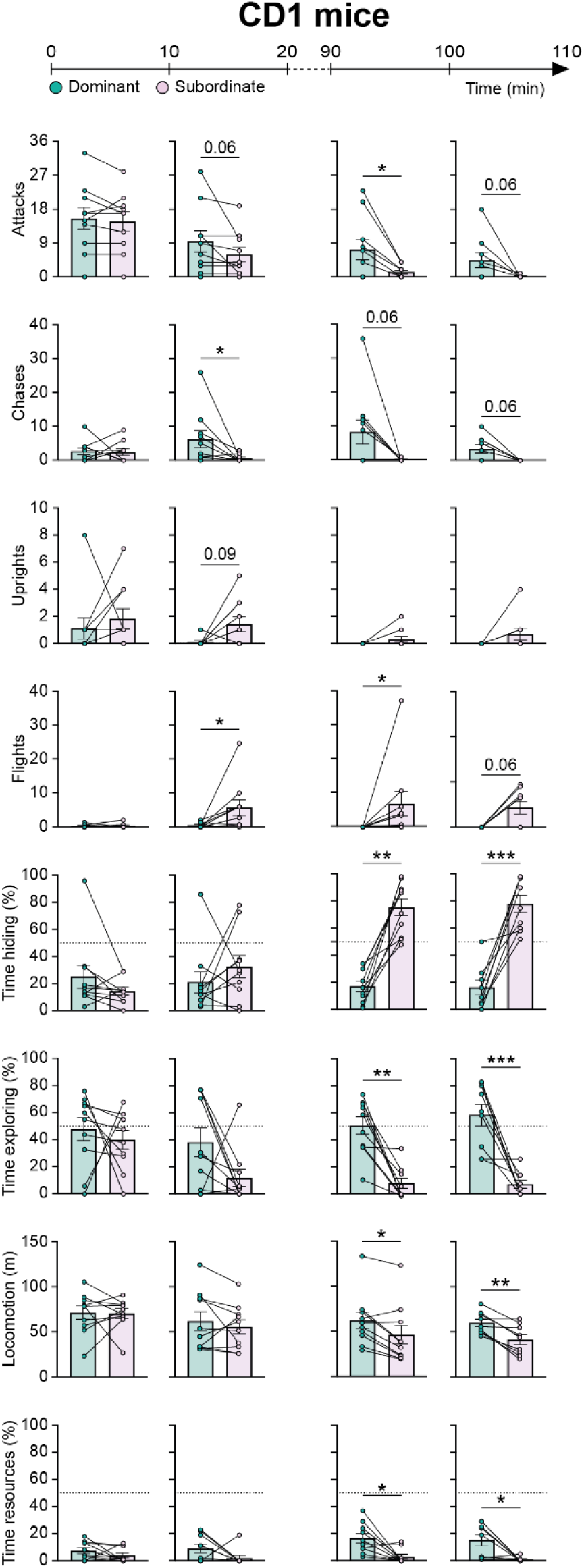
Evolution of territorial behavior in CD1 mice. Quantification of the evolution of territorial behavior differences between dominant and subordinate CD1 mice across subintervals (10 minutes) of the early and late observation periods (N = 10; Wilcoxon matched-pairs signed rank test: *P < .05, **P < .01, ***P < .001; mean ± SEM).

**Figure S2.**
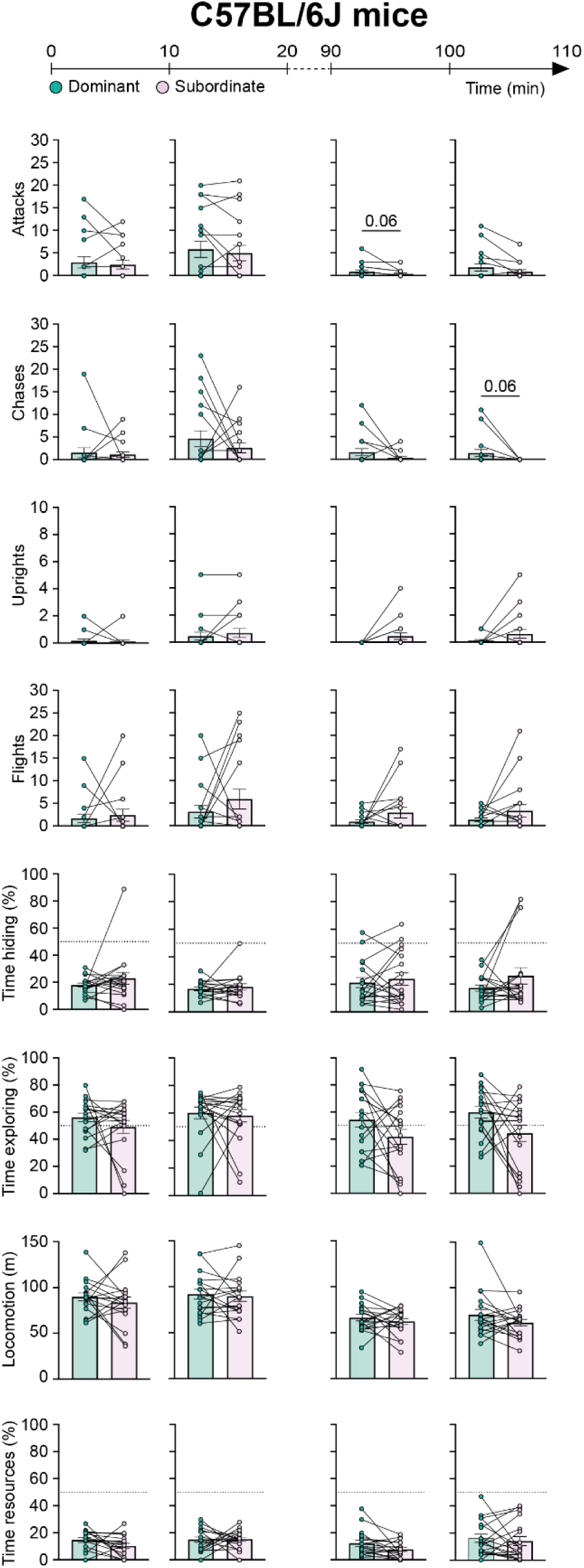
Evolution of territorial behavior in C57BL/6 mice. Quantification of the evolution of territorial behavior differences between dominant and subordinate C57BL/6 mice across subintervals (10 minutes) of the early and late observation periods (N = 18; Wilcoxon matched-pairs signed rank test: *P < .05, **P < .01, ***P < .001; mean ± SEM).

**Figure S3.**
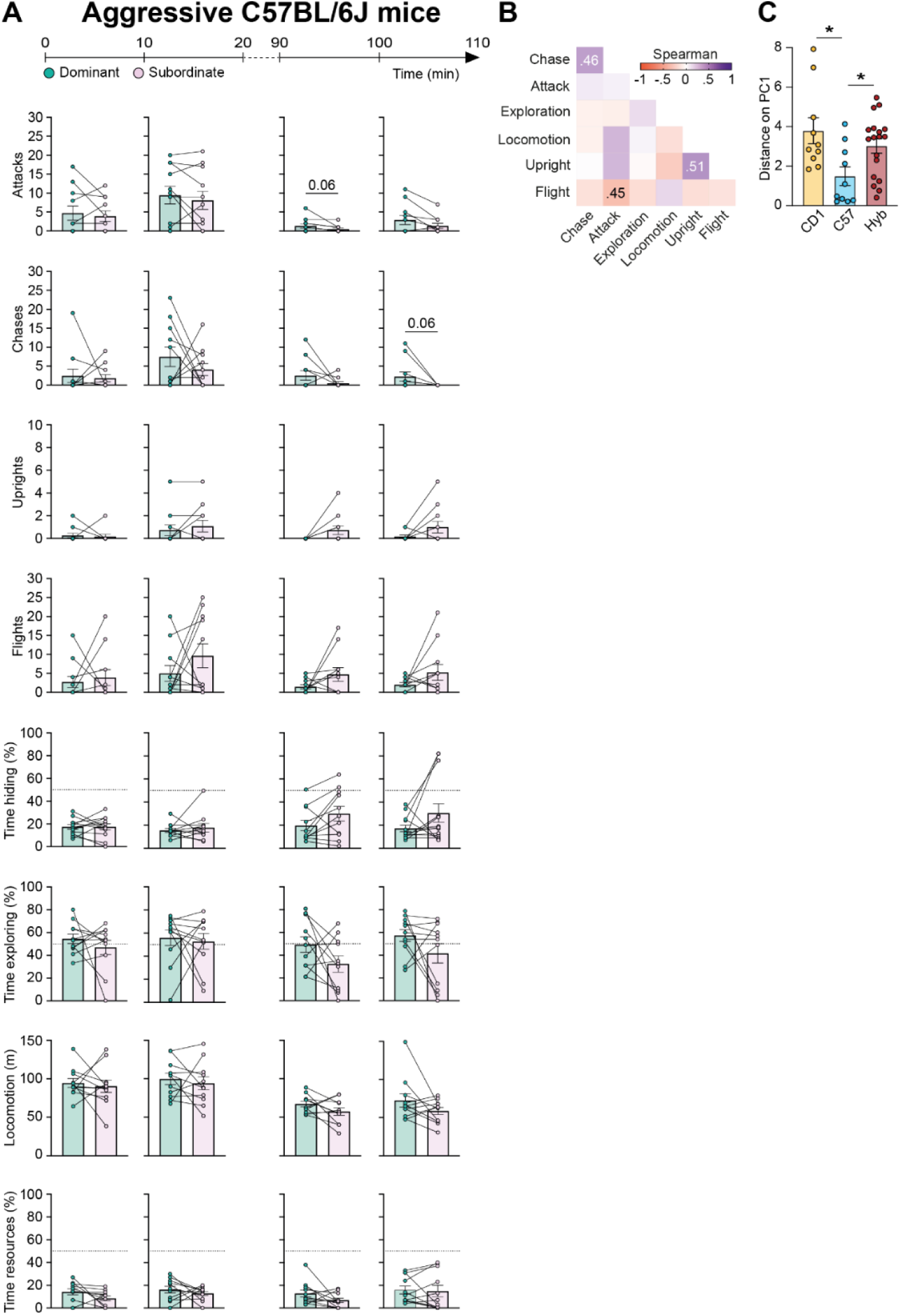
Evolution of territorial behavior in aggressive C57BL/6 mice. (**A**) Quantification of the evolution of territorial behavior differences between dominant and subordinate aggressive C57BL/6 (pairs showing at least one attack, N = 11) mice across subintervals (10 minutes) of the early and late observation periods (Wilcoxon matched-pairs signed rank test: *P < .05, **P < .01, ***P < .001; mean ± SEM). (**B**) Correlation matrix between territorial behaviors within aggressive C57BL/6 mice (N = 11 pairs, only significant correlation coefficients are reported). (**C**) Comparison of PC1 difference (dominant minus subordinate) values between strains revealed that a significant difference persisted between aggressive C57BL/6 mice and the other strains (Dunn correction for pairwise comparisons: *P < 0.01; mean ± SEM).

**Figure S4.**
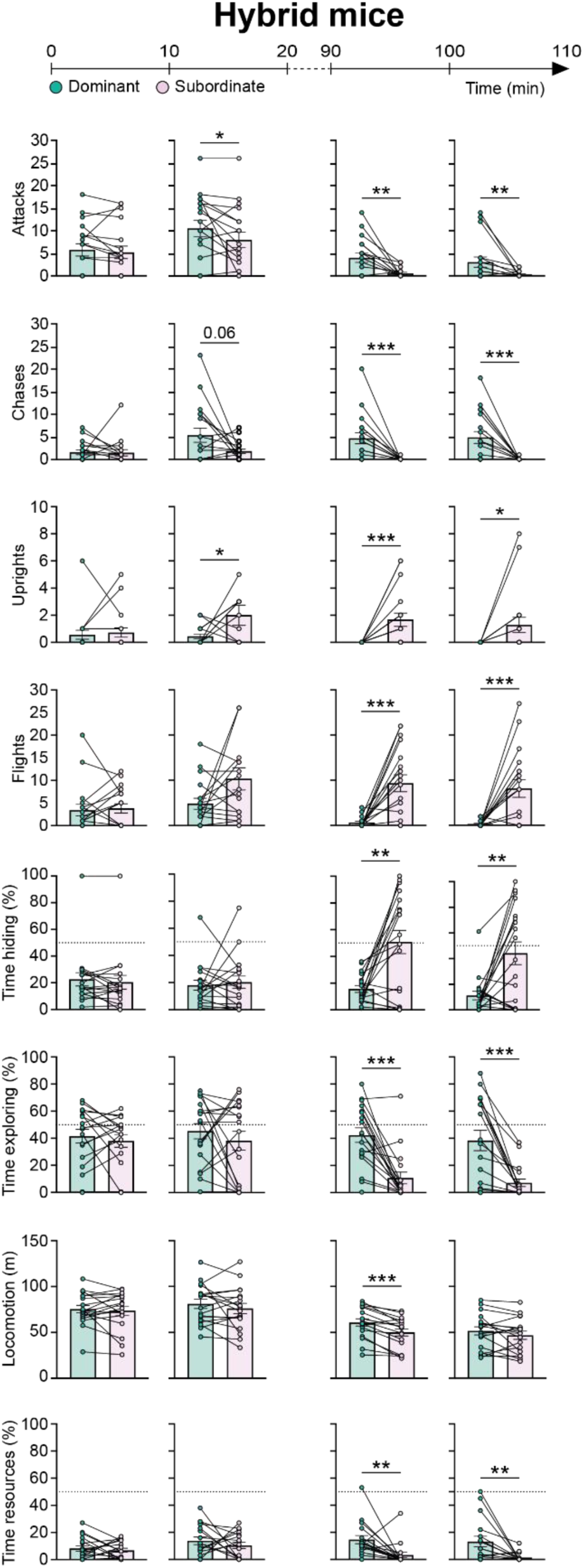
Evolution of territorial behavior in CD1xB6 F1 hybrid mice. Quantification of the evolution of territorial behavior differences between dominant and subordinate CD1xB6 F1 hybrid mice across subintervals (10 minutes) of the early and late observation periods (N = 18; Wilcoxon matched-pairs signed rank test: *P < .05, **P < .01, ***P < .001; mean ± SEM).

**Figure S5.**
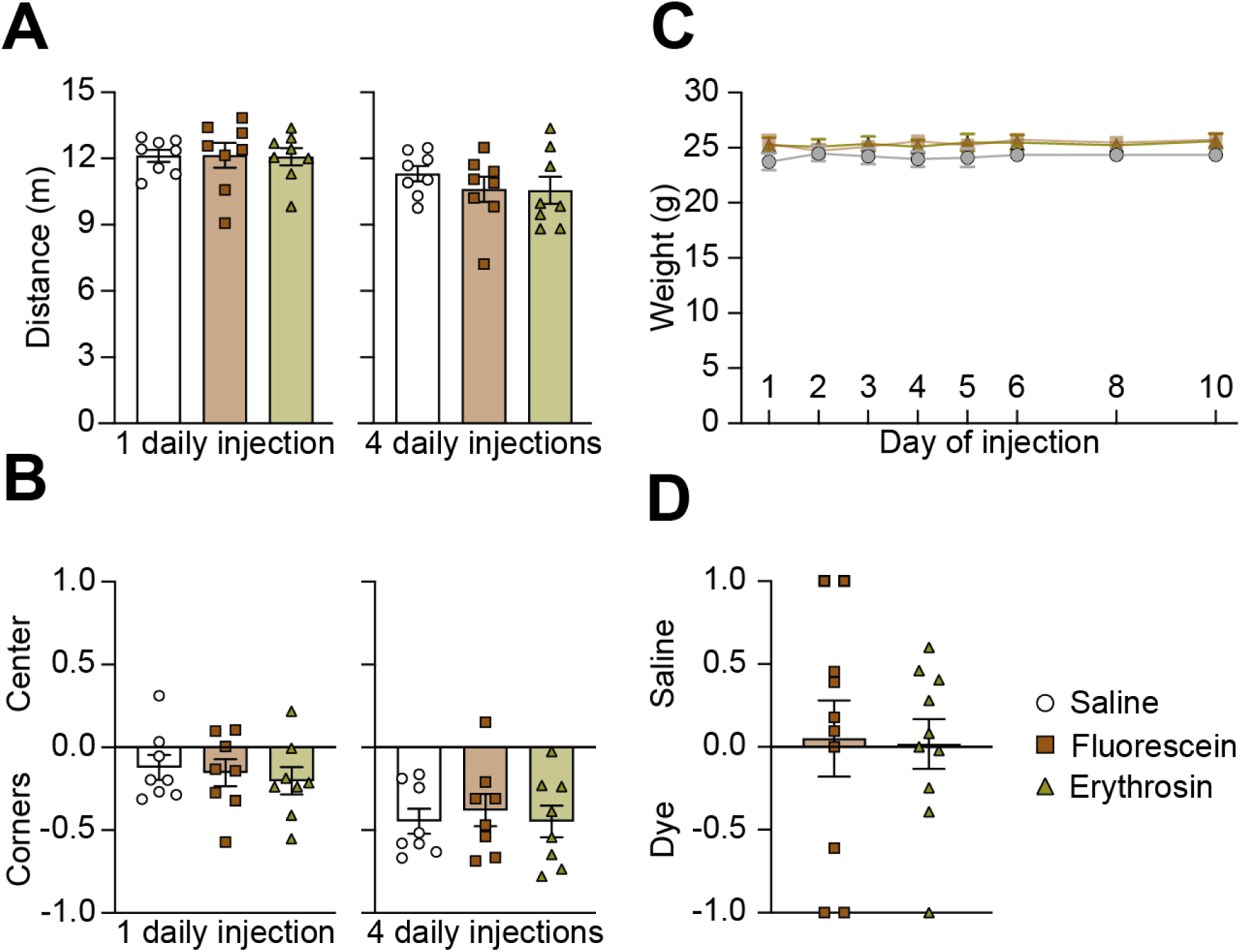
Behavioral impact of urine marking dye injection. Treatment of C57BL/6 mice with either acute or six days injection of either fluorescein or erythrosin b was not associated with significant differences in (**A**) distance traveled (left) or (**B**) time in center vs corners (right, anxiety-like behavior index: values of 1 and −1 indicate, respectively, a preference for the center or corners of the open field with an index value of 0 indicating no preference in the open field (N = 8; Dunn correction for pairwise comparisons: all P > 0.99). (**C**) Similar chronic treatment (once daily injection for ten days) of C57BL/6 mice was not associated with significant differences in body weight (N = 8). (**D**) Odor preference for urine collected from C57BL/6 mice subjected to acute treatment was not significantly different from that expressed toward urine collected from saline treated controls (N = 10; index values of −1 and 1 indicate, respectively, a preference for the dye-containing or control urine; the index value of 0 indicates no preference; Mann-Whitney *U* test: P = 0.8; mean ± SEM).

## Notes

https://git.embl.de/grp-gross/induction-of-territorial-behavior-and-dominance-hierarchies-in-laboratory-mice

### Competing Interest Statement

The authors have declared no competing interest.

